# Interneuron Control of *C. elegans* Developmental Decision-making

**DOI:** 10.1101/2021.11.07.467589

**Authors:** Cynthia M. Chai, Mahdi Torkashvand, Maedeh Seyedolmohadesin, Heenam Park, Vivek Venkatachalam, Paul W. Sternberg

**Affiliations:** Division of Biology and Biological Engineering, California Institute of Technology, Pasadena 91125, California, USA; Department of Physics, Northeastern University, Boston 02115, Massachusetts, USA

**Keywords:** Developmental plasticity, interneurons, neuropeptides, G-protein coupled receptor, pheromone, circuits, metabotropic glutamate receptor, inter-tissue signaling, physiology, metabolism

## Abstract

Animals integrate external stimuli to shape their physiological responses throughout development. In adverse environments, *Caenorhabditis elegans* larvae can enter a stress-resistant diapause state with arrested metabolism and reproductive physiology. Amphid sensory neurons feed into both rapid chemotactic and short-term foraging mode decisions, mediated by amphid and premotor interneurons, as well as the long-term diapause decision. We identify amphid interneurons that integrate pheromone cues and propagate this information via a neuropeptidergic pathways to influence larval developmental fate, bypassing the pre-motor system. AIA interneuron-derived FLP-2 neuropeptide signaling promotes reproductive growth and AIA activity is suppressed by pheromone. FLP-2 acts antagonistically to the insulin-like INS-1. FLP-2’s growth promoting effects are inhibited by upstream metabotropic glutamatergic signaling and mediated by the broadly-expressed neuropeptide receptor NPR-30. Conversely, the AIB interneurons and their neuropeptide receptor NPR-9/GALR2 promote diapause entry. These neuropeptidergic outputs allow reuse of parts of a sensory system for a decision with a distinct timescale.

## INTRODUCTION

Natural environments are highly dynamic and this complexity challenges animals to accurately integrate external cues to shape their responses. Phenotypic plasticity is the ability of a single genotype to produce more than one phenotype depending on the external environment experienced. Adaptive developmental plasticity enables organisms to remodel their physiology, morphology, and behavior to better suit the predicted future environment and ultimately enhance their ecological success (Kelly et al., 2012). Developmental plasticity often results in long-lasting changes such as switches in developmental trajectories that irreversibly alter the adult phenotype or, in some cases, induce reversible developmental arrest. This adaptive strategy is employed throughout the animal kingdom by nematodes (Bento et al., 2010; Cassada and Russell, 1975), arthropods (Brakefield et al., 1998; Laforsch et al., 2006; Moczek and Emlen, 2000), fish (Podrabsky and Hand, 1999), reptiles (Janzen and Paukstis, 1991), and amphibians (Michimae and Wakahara, 2001; Pfennig, 1992). Although it is known that sensory cues usually trigger the developmental switch and that downstream inter-tissue signaling pathways enact the alternative developmental phenotype, the integrative neural mechanisms that transduce external inputs into effector pathways are less clear (Beldade et al., 2011; Nijhout, 2003). Developmental fate choice requires sensory evidence accumulation to surpass an innately-defined threshold, signaling the appropriateness of selecting a particular trajectory over the other. Understanding how an animal generates a neural representation of current and forecasted environmental conditions and converts these circuit computations into a predictive adaptive physiological response might provide fundamental insights into the molecular and cellular basis of decision-making over developmentally-relevant timescales.

Here, we leverage the well-defined genetics and nervous system of the microscopic roundworm *Caenorhabditis elegans* to dissect the sensory integration pathways controlling larval developmental plasticity. Under adverse environmental conditions, *C. elegans* L1 stage larvae can enter an alternate stress-resistant diapause state during which metabolism and reproductive physiology are arrested(Golden and Riddle, 1984). The transition into diapause is accompanied by adaptations that enable the larva to endure and survive deleterious conditions. Dauer larvae possess thickened cuticles, enlarged intestinal lipid droplets, arrested gonad development, and are radially constricted compared to their reproductive L3 stage counterparts (Cassada and Russell, 1975). They also exhibit dauer stagespecific behaviors such as nictation, carbon dioxide attraction, and cessation of pharyngeal pumping (Cassada and Russell, 1975; Hallem et al., 2011; Lee et al., 2011). Once in the dauer stage, larvae can survive for several months without food and can revert to the reproductive trajectory at any point during this period should a favorable shift in external conditions occur (Cassada and Russell, 1975; Klass and Hirsh, 1976). The *C. elegans* larval decision to enter diapause is informed by multimodal sensory cues (Figure 1A). Coincident detection of high pheromone concentrations, high temperatures, and low food availability by the larval sensory apparatus triggers diapause entry (Golden and Riddle, 1982; 1984). *C. elegans* constitutively secretes a mixture nematode-specific glycosides called ascarosides that comprise dauer larvae-inducing pheromone, which serves as a measure of local conspecific competition for limited resources (Butcher et al., 2008; Jeong et al., 2005; Pungaliya et al., 2009). These small-molecule ascarosides bind to seven transmembrane domain receptors located in chemosensory neuron cilia to trigger intracellular signaling cascades that alter neuronal activity (Kim et al., 2009; McGrath et al., 2011).

**Figure 1.**
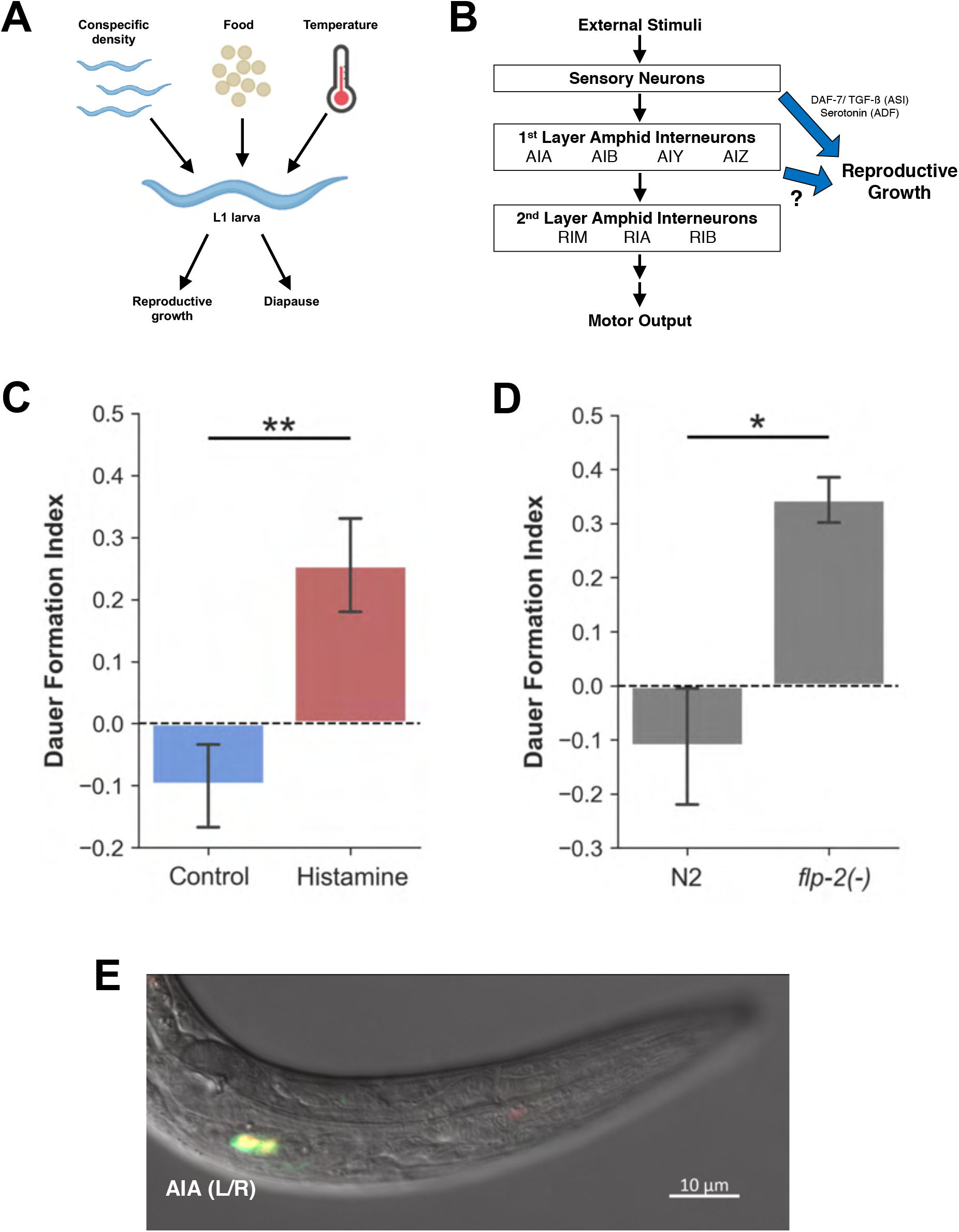
AIA inhibition recapitulates the increased dauer formation phenotype of a *flp-2* mutant. (A) Schematic illustration showing different multimodal environmental inputs integrated by an L1 larva during the diapause entry decision. (B) Schematic illustration showing hierarchical organization of neural processing layers controlling *C. elegans* behavior. (C) Dauer formation assays for *AIA::HisCl1* animals grown on plates with and without histamine. N=9 population assays. Data represented as mean ± SEM, Welch’s T-test, **p<0.01. (D) Dauer formation assays for *flp-2(-)* mutants. N=5-6 population assays. Data represented as mean ± SEM, Welch’s T-test, *p<0.05. (E) *Pflp-2::GFP* colocalization with AIA-specific *Pgcy-28.d::mCherry-H2B* marker in an L1 stage larva.

The majority of previous literature on the neural control of *C. elegans* diapause entry has focused on the contributions of sensory neurons. Systematic laser and genetic ablations of sensory neuron classes have revealed that six of the eleven amphid chemosensory neuron classes mediate diapause entry (Bargmann and Horvitz, 1991; Kim *et al*., 2009; Schackwitz et al., 1996). In *C. elegans*, sensory neurons are also the main sources of neuroendocrine ligands like DAF-7/TGF-ß (ASI neurons) and neuromodulators like serotonin (ADF neurons) that promote reproductive growth by acting on multiple tissues (Schackwitz *et al*., 1996; Sze et al., 2000). It is thus plausible that sensory layer circuits account for all integrative computations of multimodal stimuli, which then bypasses interneuron processing by directly executing secretory signaling pathways to influence developmental fate. In contrast, *C. elegans* neurobehavioral studies generally involve a hierarchical organization of information flow from the sensory layer to interneurons and finally to effector motor neurons (Figure 1B; Faumont et al., 2012). The first layer amphid interneurons AIA, AIB, AIY, and AIZ are so termed because they receive extensive synaptic inputs from sensory neurons residing within the amphid organ in the head (White et al., 1986). They then relay this information to the second layer amphid interneurons RIA, RIB, and RIM, which are presynaptic to command-like interneurons (White et al., 1986).

In this study, we combine classical genetics methods with chemogenetics, calcium imaging, and precision genome editing to examine whether a similar circuit hierarchy is involved in controlling *C. elegans* developmental plasticity (Figure 1B). To model the stressful dauer-inducing conditions encountered by *C. elegans* in the wild, we incubated developing larvae at elevated temperatures while continuously exposing them to crude pheromone extract and a low-quality, limited food source (Lee et al., 2017). The number of larvae that developed into dauers and non-dauer reproductive adults were scored and used to calculate a dauer formation index value for each assay plate (see Methods). A more positive index value indicates a greater tendency to form dauers while a more negative index value denotes a preference for entering the reproductive developmental trajectory. We find that the AIA interneurons contribute to developmental fate choice by integrating conspecific cues and propagating this information via neuropeptide signaling, in particular by the RF-amide neuropeptide encoded by *flp*-2. AIA-derived FLP-2 signaling promotes reproductive growth and is inhibited by upstream glutamatergic signaling via the metabotropic glutamate receptor MGL-1. FLP-2 signaling, which we show antagonizes insulin-like INS-1 signaling, is mediated by the downstream neuropeptide G-protein coupled receptor NPR-30, expressed in neurons and the intestine. Furthermore, AIA’s major postsynaptic partners, the NPR-9/GALR2-expressing AIB interneurons, promotes diapause entry suggesting an antagonistic relationship between both interneuron classes in this decision-making paradigm. Our findings reveal a role for the first layer amphid interneurons in coupling sensory stimuli detection to downstream neuropeptide pathways mediating inter-tissue signaling during developmental decision-making.

## RESULTS

### AIA inhibition recapitulates the increased dauer formation phenotype of a *flp-2* mutant

To investigate if AIA activity affects larval developmental trajectory, we pharmacologically silenced AIA by expressing the *Drosophila* histamine-gated chloride channel HisCl1 (Pokala et al., 2014). *C. elegans* neither synthesizes nor uses histamine as an endogenous neurotransmitter (Chase and Koelle, 2007). We constitutively silenced AIA by growing *AIA::HisCl1* animals on plates containing histamine and compared dauer formation with animals of the same genotype grown on control plates without histamine. We found that a significantly higher proportion of AIA-silenced animals entered diapause compared to the negative control, indicating that AIA promotes entry into the reproductive growth trajectory (Figure 1C).

As AIA inhibition increases dauer formation, AIA likely promotes reproductive growth. Given the relatively long timescale of the diapause entry decision, we considered the role of modulatory neuropeptides. The *flp* gene family encodes FMRFamide-like neuropeptides, some of which have been implicated in regulating pheromone-induced dauer formation (Lee *et al*., 2017). Similar to chemogenetic silencing of AIA, *flp-2(ok3351)* mutants form more dauers compared to wild-type controls (Figure 1D; Lee *et al*., 2017). The *ok3351* allele harbors a deletion that removes the first exon of both predicted *flp-2* transcripts and is thus likely to be a null mutation. *flp-2* expression has been reported in the AIA, RID, PVW, I5 and MC neurons (Kim and Li, 2004). A transcriptional reporter containing a truncated 2 kb fragment of upstream regulatory sequence 5’ to the *flp-2* start codon fused to GFP was expressed strongly and consistently in a single pair of neurons in the head during the L1 stage (Figure 1E). We identified this pair of neurons as the AIA interneurons based on GFP expression colocalization with a nuclear-localized AIA-specific mCherry marker (Figure 1E; Shinkai et al., 2011).

### AIA-derived FLP-2 promotes larval entry into the reproductive growth trajectory

To determine if FLP-2 mediates AIA’s reproductive growth-promoting role, we expressed *flp-2* cDNA specifically in AIA where it was sufficient to rescue the *flp-2(-)* dauer formation phenotype (Figure 2A). Some non-neuronal GFP expression was also visible in the intestine of the *flp-2* transcriptional GFP reporter but intestine-specific *flp-2* cDNA expression did not affect *flp-2(-)* mutant dauer formation (Figure S1). Since loss of FLP-2 signaling in the *flp-2(-)* mutant leads to increased dauer formation, we wondered if an excess of the neuropeptide would have the opposite effect on larval development. Indeed, AIA-specific overexpression of *flp-2* cDNA resulted in reduced dauer formation compared to control animals (Figure 2C).

**Figure 2.**
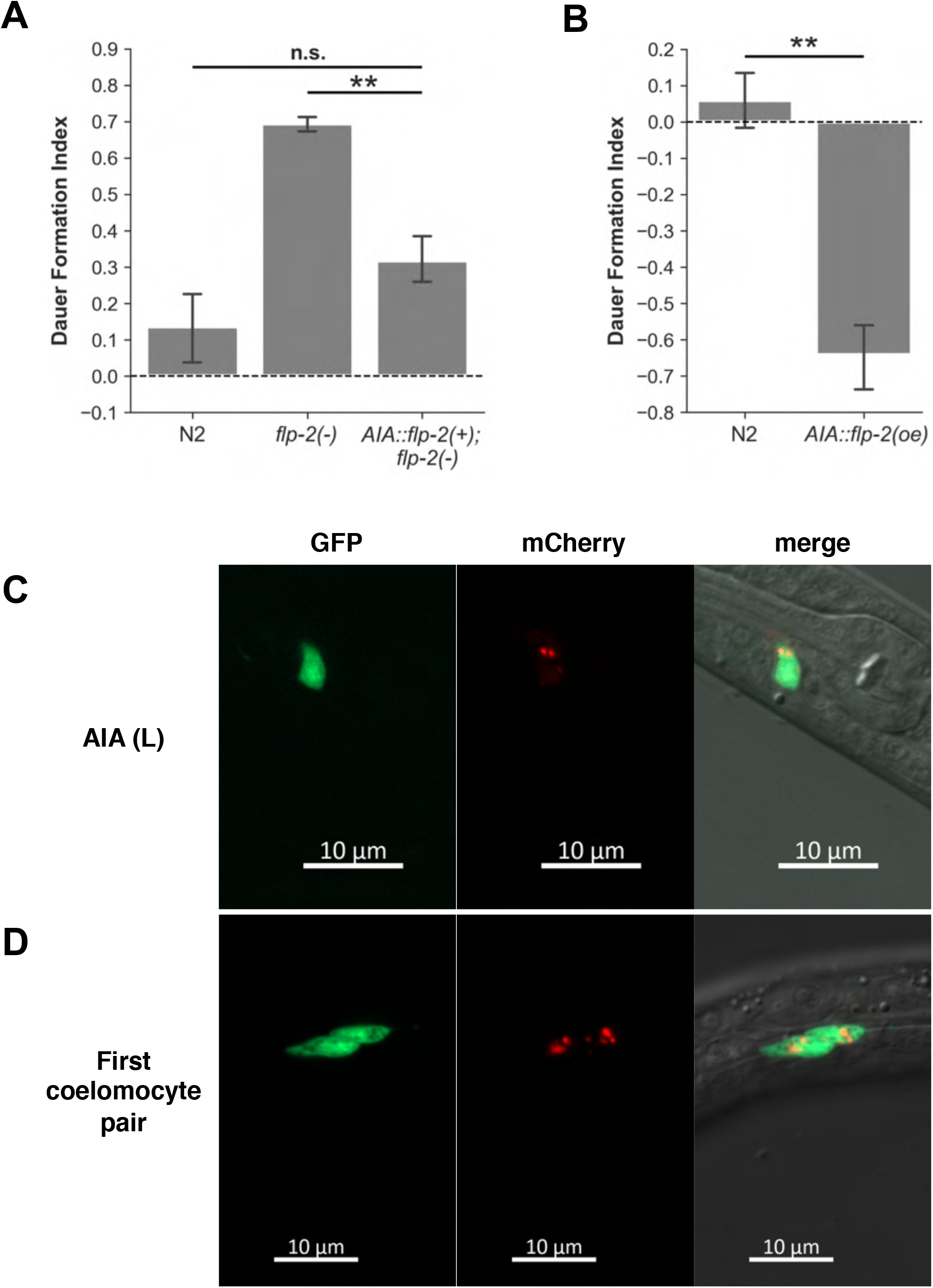
AIA-derived FLP-2 promotes larval entry into the reproductive growth trajectory. (A) Dauer formation assays for AIA-specific *flp-2* cDNA expression in an *flp-2(-)* mutant background. N=5 population assays. Data represented as mean ± SEM, ANOVA followed by Tukey HSD posthoc, **p<0.01, n.s.=no significance. (B) Dauer formation assays for AIA-specific *flp-2* cDNA overexpression. N=3 population assays. Data represented as mean ± SEM, Welch’s T-test, **p<0.01. (C) *Pflp-2::flp-2 cDNA::mCherry* colocalization with AIA-specific *Pgcy-28.d::GFP* in an L1 stage larva. (D) FLP-2::mCherry accumulation within first pair of coelomocytes visualized with *Pofm-1::GFP* in an L1 stage larva.

To confirm that FLP-2 peptides are secreted by AIA, we fused mCherry to the C-terminus of FLP-2 precursors expressed in AIA. We detected both diffuse and vesicular mCherry expression in AIA’s soma, which was visualized by soluble GFP (Figure 2D). In *C. elegans*, secreted proteins are released into the pseudocoelom and endocytosed by the coelomocyte scavenger cells (Fares and Greenwald, 2001). The six coelomocyte scavenger cells are located in three pairs along the animal’s body. During the L1 stage, we observed strong mCherry signal within GFP-expressing coelomocyte cells indicating FLP-2 peptide release and uptake (Figure 2E). Taken together, these results indicate that FLP-2 is an AIA-secreted signal that promotes reproductive development.

### Pheromone activates ASK and ADL sensory neurons but inhibits downstream AIA interneurons

The AIA interneurons make several chemical and electrical synaptic connections with the amphid chemosensory neuron classes including ASK and ADL, which mediate adult responses to certain ascarosides (Figure 3A; Jang et al., 2012; Macosko et al., 2009; White *et al*., 1986). Laser microbeam ablation and mutant analysis of ASK and ADL indicate that these sensory neurons promote dauer formation (Bargmann and Horvitz, 1991; Kim et al., 2009; Schackwitz et al., 1996). To examine the relationship between sensory neuron activity and dauer formation, we silenced ASK and ADL by selective expression of *Drosophila* HisCl1. Silencing of either neuron class reduced dauer formation suggesting that ASK and ADL activity promotes diapause entry (Figure 3B,D). We note that the effect size of ADL silencing is smaller compared to ASK silencing suggesting a subtler physiological contribution by ADL during dauer formation. In adult animals, ASK intracellular calcium levels are decreased by exposure to synthetic ascaroside cocktails that mediate adult aggregation (Macosko et al., 2009). However, the crude pheromone extract used in this study is a mixture of ascaroside components in naturally-occurring proportions whose synergistic effects might elicit different responses. We sought to determine if and how crude pheromone extract modulates ASK and ADL responses by delivering a time-locked pheromone stimulus to restrained L4 stage animals expressing the genetically-encoded calcium sensor GCaMP6s in both neuron classes (Chen et al., 2013). Pheromone delivery was accompanied by a fast, robust increase in intracellular calcium levels in ASK and ADL indicating that both sensory neuron classes detect and are activated by crude pheromone (Figure 3C,E). In addition, both paired partners of each sensory neuron class displayed symmetric responses to pheromone delivery (Figure 3C,E).

**Figure 3.**
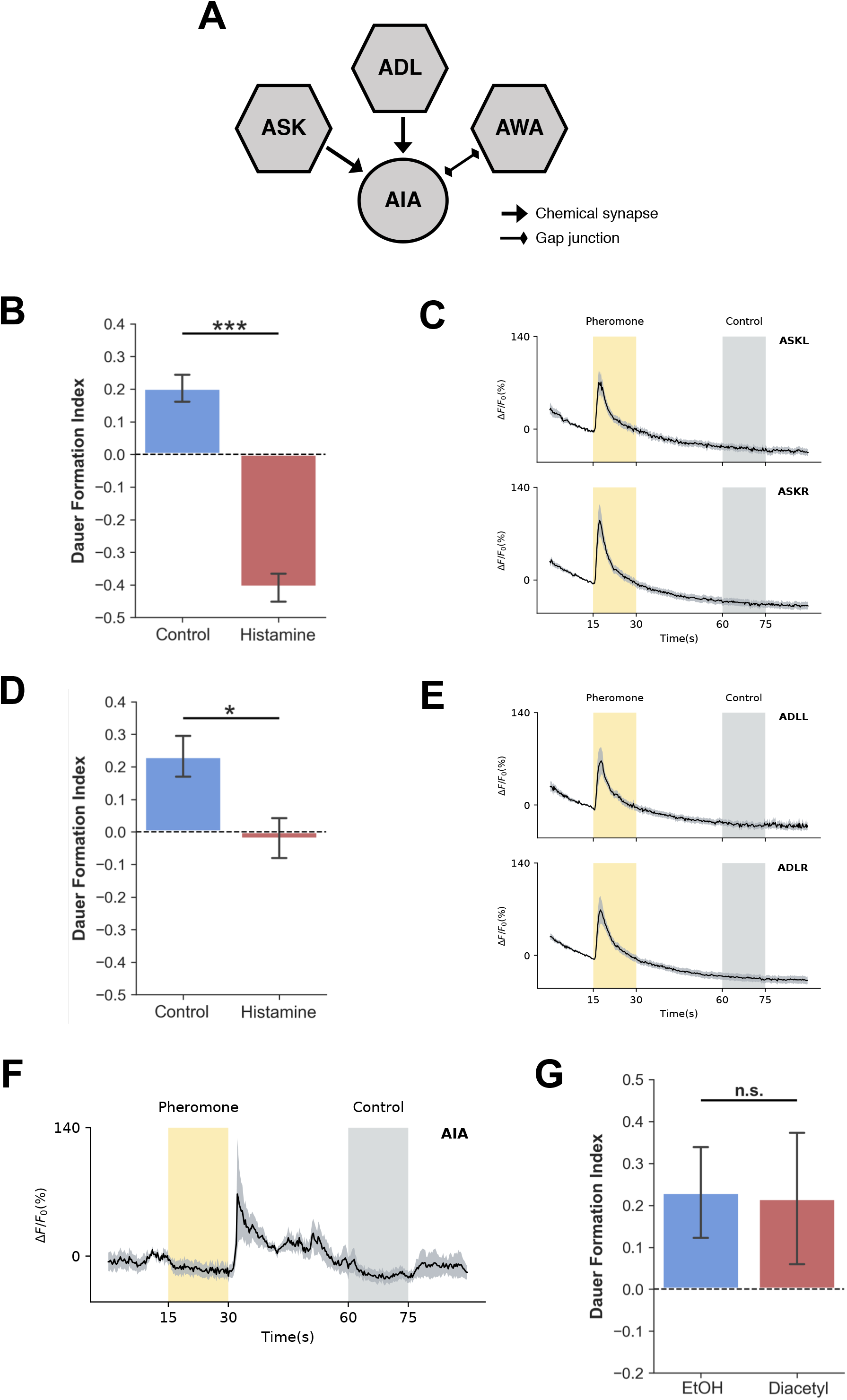
Pheromone activates ASK and ADL sensory neurons but inhibits downstream AIA interneurons. (A) Schematic illustration of unweighted synaptic connections between ASK, ADL, AWA sensory neurons and AIA. (B) Dauer formation assays for *ASK>HisCl1* animals grown on plates with and without histamine. N=7-9 population assays. Data represented as mean ± SEM, Welch’s T-test, ***p<0.001. (C) *ASK::GCaMP6s* somatic average traces in response to pheromone delivery in microfluidics device. L4 stage animals. Top panel – ASK (Left paired partner). Bottom panel – ASK (Right paired partner). N=17 animals. Data represented as mean (thick black line) ± SEM (gray shading). (D) Dauer formation assays for *ADL::HisCl1* animals grown on plates with and without histamine. N=8-11 population assays. Data represented as mean ± SEM, Welch’s T-test, *p<0.05. (E) *ADL::GCaMP6s* somatic average traces in response to pheromone delivery in microfluidics device. L4 stage animals. Top panel – ADL (Left paired partner). Bottom panel – ADL (Right paired partner). N=17 animals. Data represented as mean (thick black line) ± SEM (gray shading). (F) *AIA::GCaMP6s* process average trace in response to pheromone delivery in microfluidics device. L4 stage animals. N=5 animals. Data represented as mean (thick black line) ± SEM (gray shading). (G) Dauer formation assays for N2 animals grown in presence of 2 μL ethanol or 2 μL 1:1000 diacetyl diluted in ethanol. N=4 population assays. Data represented as mean ± SEM, Welch’s T-test, n.s.=no significance.

AIA is depolarized when animals are presented with synthetic ascaroside mixtures (Macosko et al., 2009). To probe if AIA responds similarly to crude pheromone extract, we delivered pheromone to physically-restrained L4 animals expressing GCaMP6s in AIA only. We first recorded calcium influx changes in AIA’s soma and could not detect any significant changes in AIA activity during pheromone delivery (Figure S2). As *C. elegans* interneurons exhibit compartmentalized calcium dynamics, we next measured GCaMP fluorescence changes in the AIA processes within the nerve ring where synaptic connections are densely concentrated (Hendricks et al., 2012). In our microfluidics setup, AIA is frequently quiescent with only 20% of all animals assayed displaying any process activity during experiments. In this fraction of animals, pheromone delivery consistently suppressed spontaneous AIA process activity and removal of the pheromone stimulus was followed by a post-inhibitory activity rebound that was greater in amplitude compared to before pheromone delivery (Figure 3F). Thus, AIA activity is inhibited by crude pheromone extract. Furthermore, the opposite responses to pheromone of ASK and ADL compared to AIA implies that ASK and ADL make inhibitory connections to downstream AIA interneurons.

The AWA olfactory sensory neurons sense and mediate *C. elegans* chemoattraction to the odorant diacetyl, and are gap-junctioned to AIA (Figure 3A; Bargmann et al., 1993; White et al., 1986). Diacetyl is a bacterial metabolite and might serve as a growth-promoting food signal to the bacteriovorous *C. elegans.* An AND-gate logic model of AIA integrative function has previously been proposed where coincident inhibition of glutamatergic chemosensory neurons including ASK and activation of AWA by diacetyl leads to robust AIA activation in adult animals (Dobosiewicz et al., 2019). We wondered if exposure to diacetyl concentrations that mediate adult behavioral responses might bias larval development towards the reproductive trajectory under our pheromone dauer-inducing assay conditions. However, we found that diacetyl exposure throughout the assay duration did not affect dauer formation compared to control assays with ethanol diluent only (Figure 3G). Behavioral responses to diacetyl differ between larval and adult *C. elegans*, with larvae displaying a weaker response (Fujiwara et al., 2016). Thus, stage-specific differences in sensory processing might account for the lack of developmental effects of diacetyl exposure in our dauer formation assays. Furthermore, the longer timescale of diacetyl exposure during dauer formation assays might also affect sensory neuron-to-AIA circuit dynamics differently compared to acute diacetyl presentation in a microfluidics device (Dobosiewicz et al., 2019).

### FLP-2 signaling is inhibited by upstream glutamatergic transmission via the metabotropic glutamate receptor MGL-1

Although glutamate is the major excitatory neurotransmitter in the vertebrate central nervous system, glutamate can also have inhibitory effects depending on the type of glutamate receptors present in the postsynaptic membrane. ASK and ADL are glutamatergic sensory neurons (Serrano-Saiz et al., 2013). AIA expresses both the fast ionotropic glutamate-gated chloride channel GLC-3 and the slow G-protein coupled metabotropic glutamate receptor MGL-1 (Dillon et al., 2006; Greer et al., 2008; Horoszok et al., 2001; Taylor et al., 2021). Thus, it is likely that glutamatergic transmission from upstream ASK and ADL sensory neurons affects AIA-mediated FLP-2 signaling during developmental decision-making.

To test this hypothesis, we first determined if *glc-3(-)* mutants had a dauer formation phenotype. Interestingly, *glc-3(-)* mutants exhibited a wider spectrum of developmental phenotypes compared to the binary reproductive adult or arrested dauer phenotypic outcomes observed for other strains in this study. Under pheromone dauer-inducing conditions, *glc-3(-)* mutants also produce a third developmentally-arrested partial dauer phenotype that exhibits lighter-colored bodies and is not as radially-constricted as full dauers (Vowels and Thomas, 1992). *glc-3(-)* partial dauers also display pharyngeal pumping and active locomotory behavior. The intermediate partial dauer phenotype arises from either incomplete tissue remodeling during diapause entry or an inability to maintain complete tissue remodeling once in the dauer stage. When considering both partial dauers and dauers as having entered diapause, *glc-3(-)* mutants enter diapause at levels comparable to control animals (Figure 4A). If partial dauers are considered “non-dauers” that do not meet the criteria for dauer formation, then *glc-3(-)* mutants exhibit a reduced dauer formation phenotype (Figure 4B). *flp-2(-); glc-3(-)* double mutants had an additive dauer formation phenotype significantly different from both the *flp-2(-)* and *glc-3(-)* single mutant controls (Figure 4B). Moreover, *flp-2(-); glc-3(-)* double mutants form partial dauers at a level comparable to *glc-3(-)* single mutants. Taken together, these results argue against a role for inhibitory glutamatergic transmission via GLC-3 in regulating FLP-2 signaling during dauer formation (Figure 4B).

**Figure 4.**
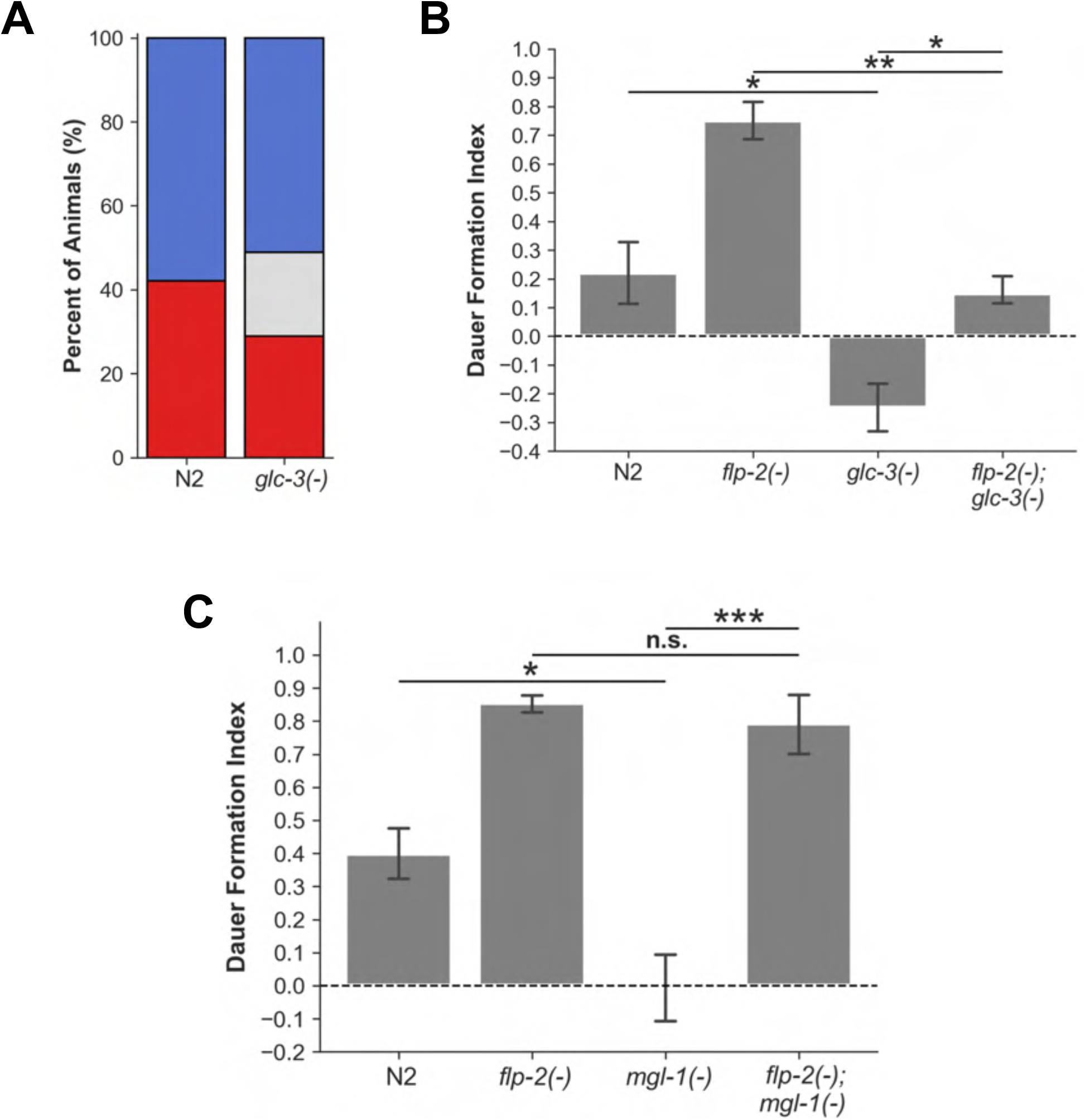
FLP-2 signaling is inhibited by upstream glutamatergic transmission via the metabotropic glutamate receptor MGL-1. (A) Percentage of full dauers (red), partial dauers (gray), and reproductive adults (blue) formed during dauer formation assays for N2 and *glc-3(-)* mutants. N=146 animals (N2) and 135 animals *(glc-3(-))*, 2 trials per genotype. (B) Dauer formation assays for *flp-2(-); glc-3(-)* double mutant genetic epistasis analysis. N=3-4 population assays. Data represented as mean ± SEM, ANOVA followed by Tukey HSD posthoc, *p<0.05, **p<0.01. (C) Dauer formation assays for *flp-2(-); mgl-1(-)* double mutant genetic epistasis analysis. N=3-4 population assays. Data represented as mean ± SEM, ANOVA followed by Tukey HSD posthoc, *p<0.05, ***p<0.001, n.s.=no significance.

In contrast, *mgl-1(-)* mutants do not form partial dauers under pheromone dauer-inducing conditions. We found that fewer *mgl-1(-)* mutants developed into dauers compared to control animals suggesting that glutamatergic transmission via MGL-1 promotes pheromone-induced dauer formation (Figure 4C, Figure S3). MGL-1 shares sequence similarity with the mammalian group II inhibitory metabotropic glutamate receptors, suggesting that ASK and ADL glutamatergic transmission might suppress AIA and FLP-2 signaling (Dillon *et al*., 2006). If FLP-2 signaling is downstream of MGL-1-mediated glutamatergic transmission, we would expect the *flp-2(-)* mutant’s increase in dauer formation to mask the *mgl-1(-)* mutant’s reduction in dauer formation phenotype in the double mutant, displaying epistasis. As *mgl-1* and *flp-2* genes are on the same chromosome, we generated a *flp-2(-); mgl-1(-)* double mutant by CRISPR mutagenesis of the *mgl-1* locus in a *flp-2(-)* mutant background (see Methods). The *flp-2(-); mgl-1(-)* double mutant phenocopied the *flp-2(-)* single mutant’s increased dauer formation phenotype indicating that *flp-2* is in the same genetic pathway as *mgl-1* and acts downstream (Figure 4C). Our results suggest that FLP-2 signaling is inhibited by glutamatergic transmission via the metabotropic glutamate receptor MGL-1.

### The broadly-expressed neuropeptide receptor NPR-30 is downstream of FLP-2 signaling

We next focused on identifying downstream elements of the AIA-mediated developmental decision-making circuit. *In vitro* studies indicate that peptides cleaved from the FLP-2 precursor protein bind to the G-protein coupled neuropeptide receptor FRPR-18 in a heterologous mammalian cell expression system (Larsen et al., 2013). However, *frpr-18(-)* mutants had a decreased dauer formation phenotype (Figure 5A). Furthermore, *flp-2(-); frpr-18(-)* double mutants exhibit an additive phenotype that is significantly different from both the *flp-2(-)* and *frpr-18(-)* single mutant controls (Figure 5A). These results indicate that *flp-2* and *frpr-18* are not in the same genetic pathway in the context of dauer formation. We continued our search for potential downstream receptor mediators of FLP-2 signaling by testing neuropeptide G-protein coupled receptor mutants that have an increased dauer formation phenotype for their ability to completely suppress the *flp-2* overexpression reduced dauer formation phenotype; this screen identified *npr-30* (Figure 5B). Thus, *npr-30* is downstream of *flp-2* in the same pathway mediating the diapause entry decision.

**Figure 5.**
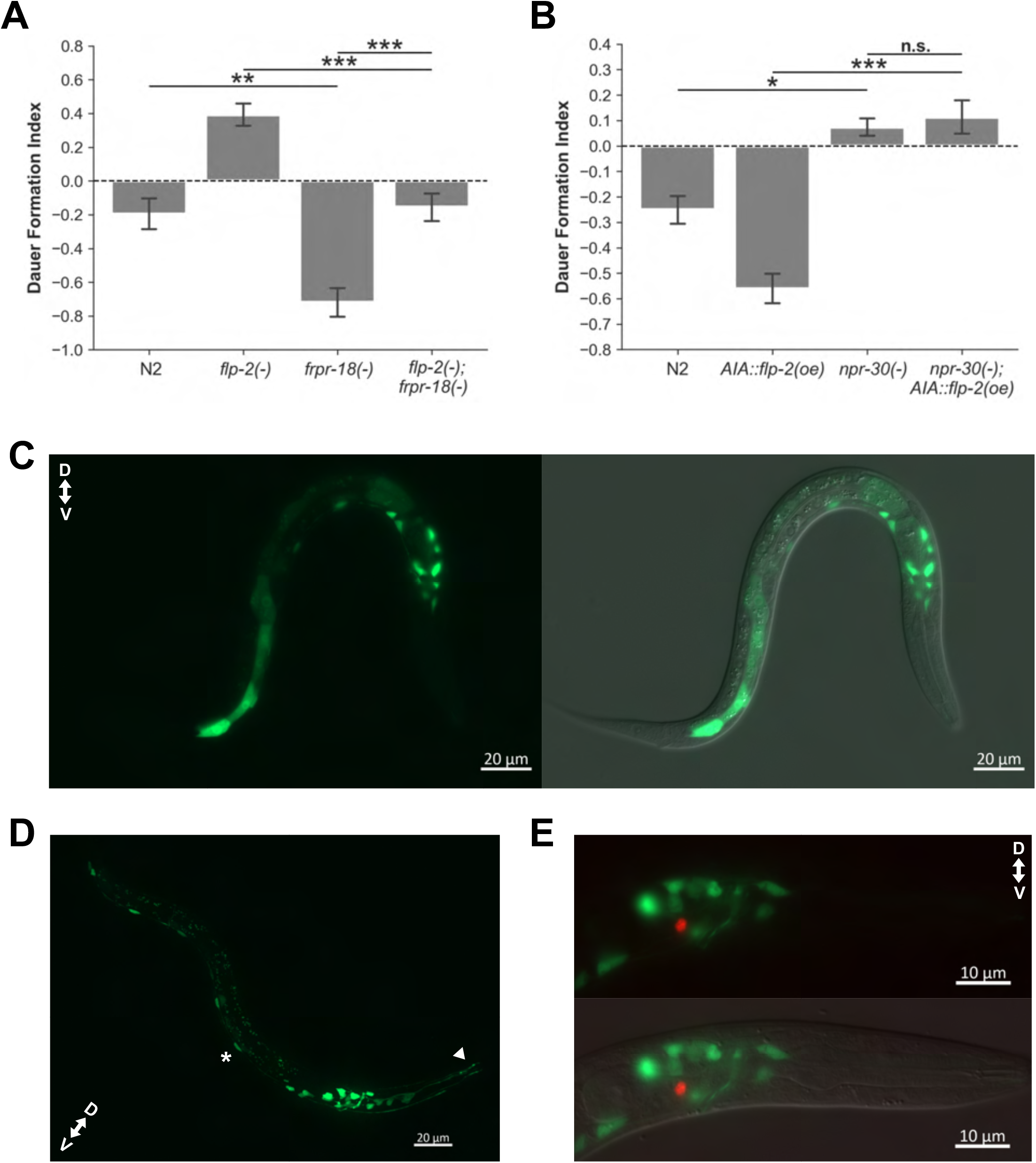
The broadly-expressed neuropeptide receptor NPR-30 is downstream of FLP-2 signaling. (A) Dauer formation assays for *flp-2(-); frpr-18(-)* double mutant genetic epistasis analysis. N=6-7 population assays. Data represented as mean ± SEM, ANOVA followed by Tukey HSD posthoc, **p<0.01, ***p<0.001. (B) Dauer formation assays for *npr-30(-)* suppression of *AIA::flp-2* cDNA overexpression. Only transgenic animals counted. N=4-8 population assays. Data represented as mean ± SEM, ANOVA followed by Tukey HSD posthoc, *p<0.05, ***p<0.001, n.s.=no significance. (C) Representative image of *npr-30* translational GFP reporter (PS9109) expression in an L1 stage larva. GFP expression present in nervous system and intestine. Left panel – GFP channel only. Right panel – GFP and DIC channels merge. (D) Representative image of *npr-30* translational GFP reporter (PS9152) expression in an L1 stage larva. Strain injected with lower GFP reporter plasmid concentration than (C) to enable better visualization of motor neurons located along the ventral side of the animal’s body. GFP is expressed in sensory neurons (white arrowhead pointing at dendritic processes extending towards nose tip), interneurons, and motor neurons (white asterisk). (E) Representative image showing AIA-specific *Pgcy-28.d::mCherry-H2B* does not colocalize with an *npr-30* translational GFP reporter in an L1 stage larva. Top panel – GFP and mCherry channels merge. Bottom panel – GFP, mCherry, and DIC channels merge.

To determine where *npr-30* is expressed, we generated a *npr-30* translational reporter containing an 8.5 kb *npr-30* genomic fragment linked to GFP by an SL2-spliced intercistronic region. During the L1 stage, GFP was broadly expressed in the nervous system as well as in the intestine (Figure 5C). In the nervous system, we observed expression in sensory neurons, interneurons, and motor neurons, suggesting that FLP-2 signaling exerts its global modulatory effects via NPR-30 (Figure 5D). Besides acting on other targets, FLP-2 might also mediate an AIA autoregulatory positive feedback loop to drive and sustain commitment to reproductive growth under favorable conditions. However, our *npr-30* translational GFP reporter did not colocalize with an AIA-specific mCherry marker in L1 larvae, suggesting this motif is unlikely (Figure 5E).

### Insulin-like growth factor INS-1 and FLP-2 have antagonistic roles in developmental decisionmaking

The insulin signaling pathway is an evolutionarily-conserved regulator of energy homeostasis and metabolism. In mammals, insulin signaling promotes lipid storage and inhibits the breakdown of fat (Huang et al., 2020). In *C. elegans*, the larval transition into diapause entails increased lipid storage and decreased fat metabolism to facilitate long-term survival under resource-scarce conditions (Ashrafi, 2007). In addition to *flp-2*, AIA co-expresses the insulin-related growth factor-encoding *ins-1* (Kodama et al., 2006). In adult animals, AIA-specific INS-1 release mediates normal turning behavior during odor-evoked local search (Chalasani et al., 2010). Therefore, we decided to investigate if AIA-derived INS-1 might play a role in dauer formation. Under pheromone dauer-inducing conditions, *ins-1(-)* mutants exhibited a decreased dauer formation phenotype indicating that INS-1 signaling promotes diapause entry (Figure 6A). Furthermore, the *flp-2(-); ins-1(-)* double mutant displayed an additive phenotype that was significantly different from both the *flp-2(-)* and *ins-1(-)* single mutant phenotypes, suggesting that these neuropeptides act in independent genetic pathways (Figure 6A). To determine if AIA-derived INS-1 promotes dauer entry, we expressed *ins-1* cDNA in an AIA-specific manner in an *ins-1(-)* mutant background and found that it did not affect the *ins-1(-)* mutant’s dauer formation phenotype (Figure 6B). As *ins-1* is expressed in many other neuron classes besides AIA, we propose that INS-1 secretion by other neuron classes antagonizes AIA-derived FLP-2 signaling during pheromone-induced dauer formation (Figure 6C; Kodama *et al*., 2006).

**Figure 6.**
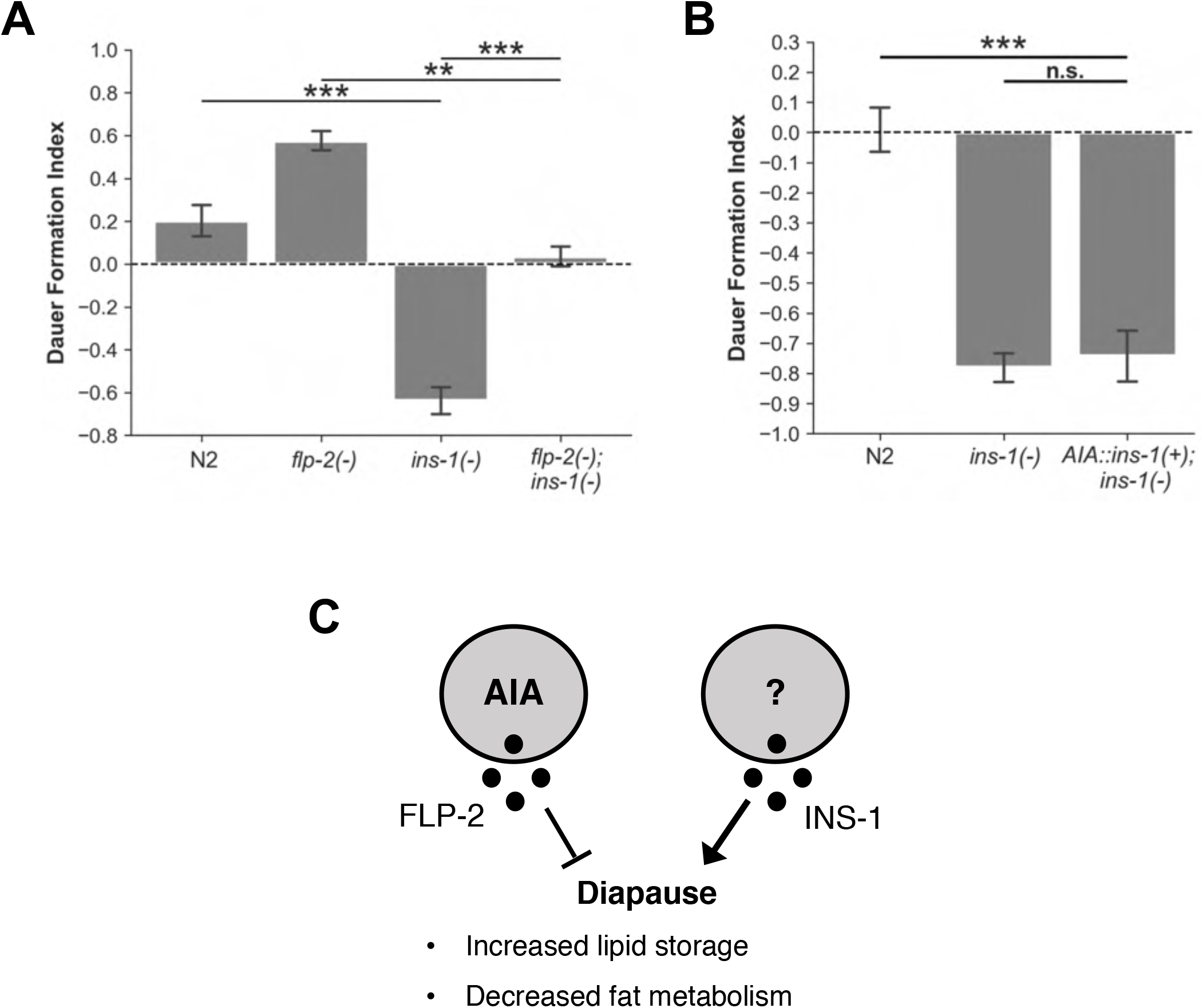
Insulin-like growth factor INS-1 and FLP-2 have antagonistic roles in developmental decision-making. (A) Dauer formation assays for *flp-2(-); ins-1(-)* double mutant genetic epistasis analysis. N=3-4 population assays. Data represented as mean ± SEM, ANOVA followed by Tukey HSD posthoc, **p<0.01, ***p<0.001. (B) Dauer formation assays for AIA-specific *ins-1* cDNA expression in an *ins-1(-)* mutant background. N=4 population assays. Data represented as mean ± SEM, ANOVA followed by Tukey HSD posthoc, ***p<0.001, n.s.=no significance. (C) Schematic model showing antagonistic relationship between FLP-2 and INS-1 signaling during diapause entry.

### AIB interneurons and their neuropeptide receptor NPR-9/GALR2 promote diapause entry

The AIB interneurons are AIA’s main postsynaptic partners and the strong unidirectional synaptic connection from AIA to AIB is likely to be inhibitory based on their opposite functions in behavioral studies (Wakabayashi et al., 2004). Furthermore, AIA is cholinergic and AIB expresses the ACC-1 acetylcholine-gated chloride channel (Pereira et al., 2015; Putrenko et al., 2005; Taylor et al., 2021). However, AIB also expresses excitatory glutamate receptors and receives synaptic inputs from glutamatergic sensory neurons including the pheromone-sensing ADL (Brockie et al., 2001; Serrano-Saiz et al., 2013; White et al., 1986). This observation suggests that excitatory glutamatergic and inhibitory cholinergic neurotransmission from the sensory layer and other pre-motor interneurons, respectively, might compete to shape AIB responses under different environmental conditions.

Pharmacological silencing of AIB reduced dauer formation suggesting that the AIA and AIB first layer interneurons have antagonistic effects on larval developmental choice (Figure 7A). *npr-9* encodes a neuropeptide G-protein coupled receptor that is homologous to human galanin receptor 2 (GALR2) and is expressed in the AIB neurons where it mediates adult foraging behavior (Bendena et al., 2008). *npr-9(-)* mutants assayed for pheromone-induced dauer formation recapitulated the AIB-silenced decreased dauer formation phenotype suggesting that AIB’s diapause entry-promoting role involves

**Figure 7.**
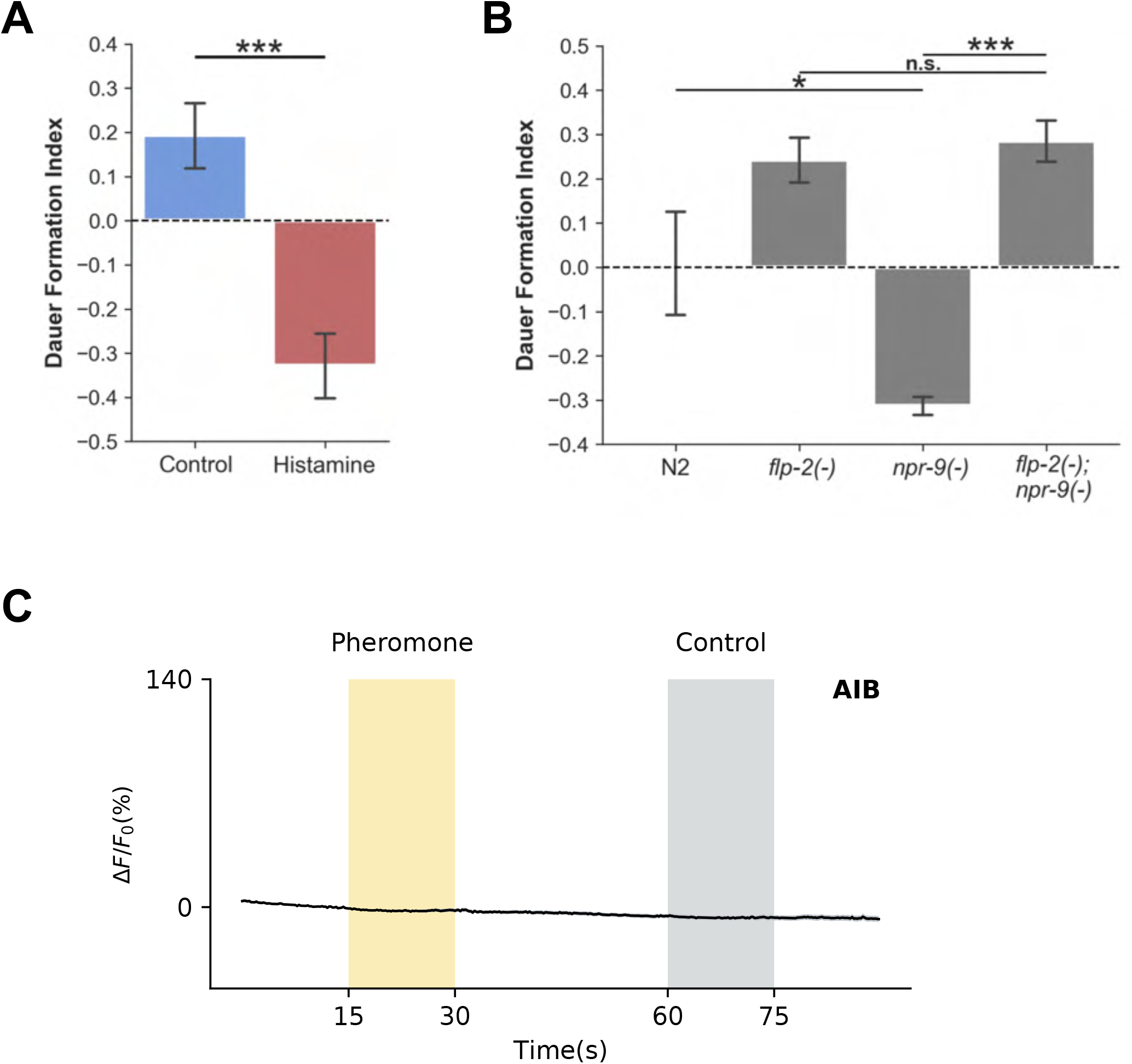
AIB interneurons and their neuropeptide receptor NPR-9/GALR2 promote diapause entry. (A) Dauer formation assays for AIB> *HisCl1* animals grown on plates with and without histamine. N=7 population assays. Data represented as mean ± SEM, Welch’s T-test, ***p<0.001. (B) Dauer formation assays for *flp-2(-); npr-9(-)* double mutant genetic epistasis analysis. N=2-4 population assays. Data represented as mean ± SEM, ANOVA followed by Tukey HSD posthoc, *p<0.05, ***p<0.001, n.s.=no significance. (C) *AIB::GCaMP6s* process average trace in response to pheromone delivery in microfluidics device. L4 stage animals. N=13 animals. Data represented as mean (thick black line) ± SEM (gray shading).

NPR-9 signaling (Figure 7B). We used a genetic epistasis test to determine whether FLP-2 and NPR-9 mediate related or independent signaling pathways. As *npr-9* and *flp-2* are also on the same chromosome, we used a similar CRISPR-based strategy as with the *flp-2(-); mgl-1(-)* strain discussed above to generate a *flp-2(-); npr-9(-)* double mutant. The *flp-2(-); npr-9(-)* double mutant phenocopied the *flp-2(-)* single mutant’s increased dauer formation phenotype indicating that *flp-2* is downstream of *npr-9* in the same genetic pathway that influences developmental fate (Figure 7B). The decreased dauer formation phenotype of AIB-silenced animals suggests that AIB is activated by adverse environmental cues that promote dauer formation. However, neither AIB’s process nor soma exhibited any activity changes in response to acute pheromone delivery in a microfluidics device (Figure 7C, Figure S4). The absence of any AIB response to pheromone presentation alone suggests that AIB activity might be gated by additional sensory inputs that could interact with the pheromone stimulus. As L4 stage animals were used for calcium imaging experiments, it is also possible that stage-specific synaptic connectivity differences might underlie the observed discrepancies between dauer formation assay results and functional imaging data.

## DISCUSSION

Here, we show that a pair of interneurons controls the switch between different larval developmental trajectories depending on prevailing environmental conditions. Under favorable conditions, AIA is active and secretes FLP-2 neuropeptides, which promotes reproductive growth (Figure 8A). FLP-2 signaling is mediated by the neuropeptide receptor NPR-30 which is broadly expressed in neurons and the intestine (Figure 8A). As conspecific population density and competition for limited resources increases, the concentration of secreted pheromone in the local area also increases. These pheromone cues activate the ASK and ADL sensory neurons, which suppress downstream AIA interneuron activity (Figure 8B). FLP-2 signaling is inhibited by upstream glutamatergic transmission mediated by the metabotropic glutamate receptor MGL-1 resulting in more larvae choosing to enter diapause (Figure 8B). Thus, we have discovered how salient sensory cues are transduced by an integrative interneuron into neuropeptidergic signals that shape larval developmental decision-making. We further expand this decision-making circuit by functionally implicating the AIB interneurons, AIA’s main postsynaptic partners, and their NPR-9/GALR2 neuropeptide receptor in promoting diapause entry (Figure 8C).

**Figure 8:**
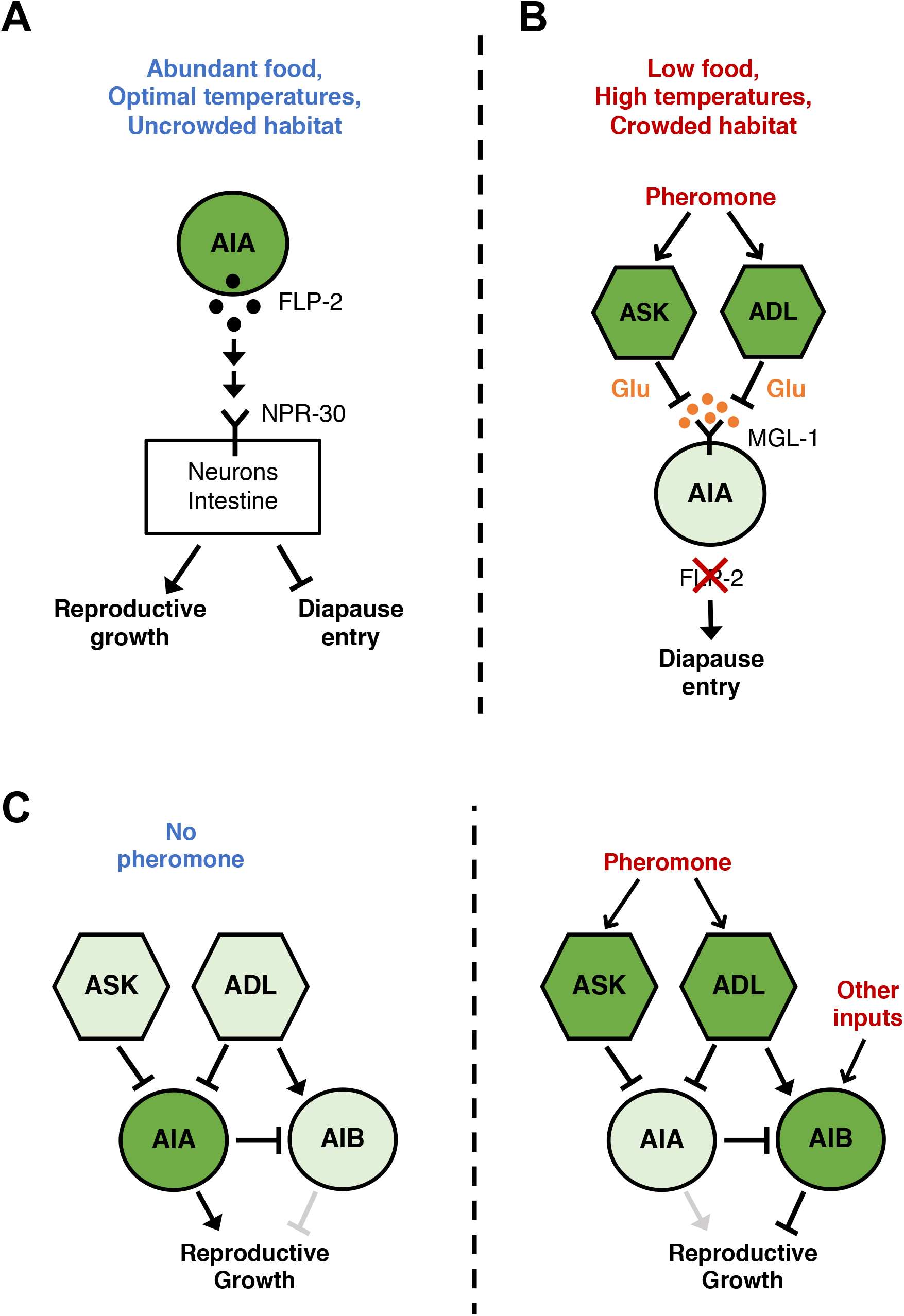
AIA controls a neuropeptide switch between different larval developmental trajectories. (A and B) Schematic model of AIA-derived FLP-2 signaling pathway under different environmental conditions: (A) Reproductive growth conditions and (B) Dauer-inducing conditions. (C) Schematic model of neural circuit mediating the diapause entry decision including pheromone-sensing neurons and interneurons identified in this study.

Diapause entry can be maladaptive if an incorrect assessment of environmental conditions is made since dauer larvae can take up to 20 hours to fully re-enter the trajectory leading to reproductive adulthood (Cassada and Russell, 1975). The metabolic costs of physiological remodeling coupled with the delay in nutritional intake (dauers are non-feeding) can negatively impact future reproductive success. Thus, we would expect such an important calculation to be under the control of complex, distributed regulatory networks as opposed to relying on a more efficient sparse-coding scheme that is less robust when challenged with noisy inputs from nature. There are likely to be more neuron classes involved in this circuit, especially other interneurons that lie at the confluence of multiple upstream sensory modalities and have neuromodulatory capabilities. A key motivation behind this study was to determine if information flow during developmental decision-making followed the same generalized hierarchical structure as behavior. Based on our findings, the interneuron layer does contribute to the diapause entry decision in an integrative capacity; it would be interesting to see whether interneuron-derived signals eventually feed back to the sensory layer, which then executes terminal effector pathways that ‘locks in’ larval fate.

The results of our microfluidics experiments show that acute inhibition of AIA by pheromone is followed by a sharp increase in calcium influx immediately after pheromone delivery is terminated. This observed post-inhibitory increase in AIA activity appears to contradict our functional dauer formation assay results which suggest that dauer-inducing environmental conditions inhibit AIA-derived FLP-2 signaling. However, the AIA response to acute pheromone presentation is likely to be mediated by the fast ionotropic glutamate-gated GLC-3 chloride channel. Based on our epistasis analysis results, FLP-2 signaling is largely mediated by the slower-acting metabotropic MGL-1 receptor. Larvae are also exposed to lower pheromone concentrations over longer timescales during dauer formation assays compared to the microfluidics setup. Furthermore, larvae encounter other multimodal sensory cues such as temperature and food during dauer formation assays, which might interact with and modulate the neuronal response to pheromone. Thus, post-inhibitory activity rebounds driven by acute neuronal inhibition might not actually be occurring during the actual diapause entry decision-making process. While acute pheromone delivery is useful for identifying the valence of a neuron’s response to the stimulus, it is clearly insufficient to accurately determine larval neuronal dynamics over developmentally-relevant timescales. Advances in long-term functional imaging of neurons in larvae cultured in agarose microchambers might well be useful for this application (Bringmann, 2011; Turek et al., 2015).

Neuropeptide signaling is an ancient inter-tissue communication mechanism that regulates animal behavioral and physiological states (Elphick et al., 2018). In *C. elegans*, individual neuron classes and non-neuronal tissue types express a constellation of neuropeptides and neuropeptide receptors underscoring the vastness and complexity of extrasynaptic neurohormonal networks present in a simple organism (Taylor et al., 2021). The arrested dauer organismal phenotype arises from coordinated multi-tissue remodeling driven by individual fate decisions computed at the cellular level. In this study, we discovered that developmentally-relevant sensory inputs likely spatially bifurcate downstream of AIA-mediated FLP-2 signaling at the NPR-30 neuropeptide receptor, which is expressed in the intestine in addition to other neuron classes and thus has the hallmarks of a fan-out circuit motif. Future studies comparing the roles of NPR-30-mediated signaling between neuronal and intestinal cell types during diapause entry might yield insights into the mechanisms of tissue-specific neuropeptide modulation. Insulin/insulin-like growth factor (IGF) signaling is a similarly conserved inter-tissue communication pathway (Garofalo, 2002; Murphy and Hu, 2013). The DAF-2 receptor tyrosine kinase is the only insulin/IGF receptor encoded by the *C. elegans* genome and is expressed in the intestine and nervous system (Hunt-Newbury et al., 2007; Kimura et al., 2011; Kimura et al., 1997). *daf-2* loss-of-function mutants have an increased dauer formation phenotype indicating that the INS-1 ligand antagonizes DAF-2 receptor signaling (Kimura et al., 1997). We have shown that FLP-2 and INS-1 mediate antagonistic signaling pathways during pheromone-induced diapause entry, and they likely converge on intestinal cells that co-express NPR-30 and DAF-2. This discovery is concordant with the intestine being a key target tissue in the diapause entry decision, and we have demonstrated a neural pathway from sensory transduction to this target tissue (Hung et al., 2014).

Obesity is an increasingly prevalent disease characterized by excessive levels of body adiposity and is itself a risk factor for other debilitating medical conditions such as cancer, diabetes, and heart disease (Blüher, 2019). G-protein coupled receptors are important regulators of metabolism and energy homeostasis, and mutations in genes encoding these metabotropic receptors are associated with altered risk for developing obesity (Akbari et al., 2021; Oliveira de Souza et al., 2021). It is striking that in the neural pathway regulating *C. elegans* larval developmental plasticity identified here, all receptors involved are G-protein coupled metabotropic receptors (MGL-1, NPR-30, NPR-9). This highlights a conserved role for metabotropic receptors in establishing and maintaining chronic metabolic and physiological states, the dysfunction of which can have profound effects on an organism’s life history.

The metabotropic glutamate receptor MGL-1 mediates foraging behavior and reproductive plasticity in adult *C. elegans*, and is expressed in just four interneuron classes AIA, RMD, AIY, and NSM (Jeong and Paik, 2017; López-Cruz et al., 2019). Here, we show that MGL-1 plays a critical role in establishing an alternative larval phenotype over developmentally-relevant timescales by inhibiting growth-promoting FLP-2 neuropeptide signaling. MGL-1 is similar in sequence to the mammalian group II glutamate metabotropic receptors mGluR2 and mGluR3, which are expressed in the hypothalamus and pituitary gland (Durand et al., 2008). When activated by glutamate binding, the group II mGluRs inhibit cyclic AMP production as a result of their negative coupling to adenylyl cyclase resulting in inhibition of the receptor-expressing neuron (Cartmell and Schoepp, 2000). In mammalian nervous systems, the hypothalamus receives inputs from multiple sensory modalities and integrates this wide range of information to drive appropriate behavioral and physiological responses (Saper and Lowell, 2014). In particular, the hypothalamic-pituitary axis of transmission mediates many aspects of growth, development, and the response to stressful stimuli through neuroendocrine regulation (Saper and Lowell, 2014). In these brain regions, glutamatergic neurotransmission via group II mGluRs inhibits the release of neuroendocrine factors (Durand et al., 2008; Scaccianoce et al., 2003). Similarly in *C. elegans*, we find that the AIA interneurons act as an integrative hub that translates inhibitory glutamatergic neurotransmission from the sensory layer into downregulation of neuropeptidergic signaling that ultimately impacts energy metabolism, reproductive physiology, and organismal morphology.

## ACKNOWLEDGMENTS

Some strains were provided by the CGC, which is funded by NIH Office of Research Infrastructure Programs (P40 OD010440). WormBase provided key molecular and genetic information. Some strains were provided by the lab of Dr. Shohei Mitani as part of the National Bioresource Project. Figure 1A was created with BioRender.com.

## AUTHOR CONTRIBUTIONS

Conceptualization – CMC, PWS. Methodology – CMC. Crude pheromone extraction and dauer formation assays – CMC. Molecular cloning and transgenesis – CMC. Microscopy – CMC. CRISPR mutagenesis – HP. Calcium imaging – MT, MS. Data analysis – CMC, MT, MS. CMC wrote the paper with input from all authors. Funding acquisition – PWS, VV.

---

**Supplementary Figure 1:**
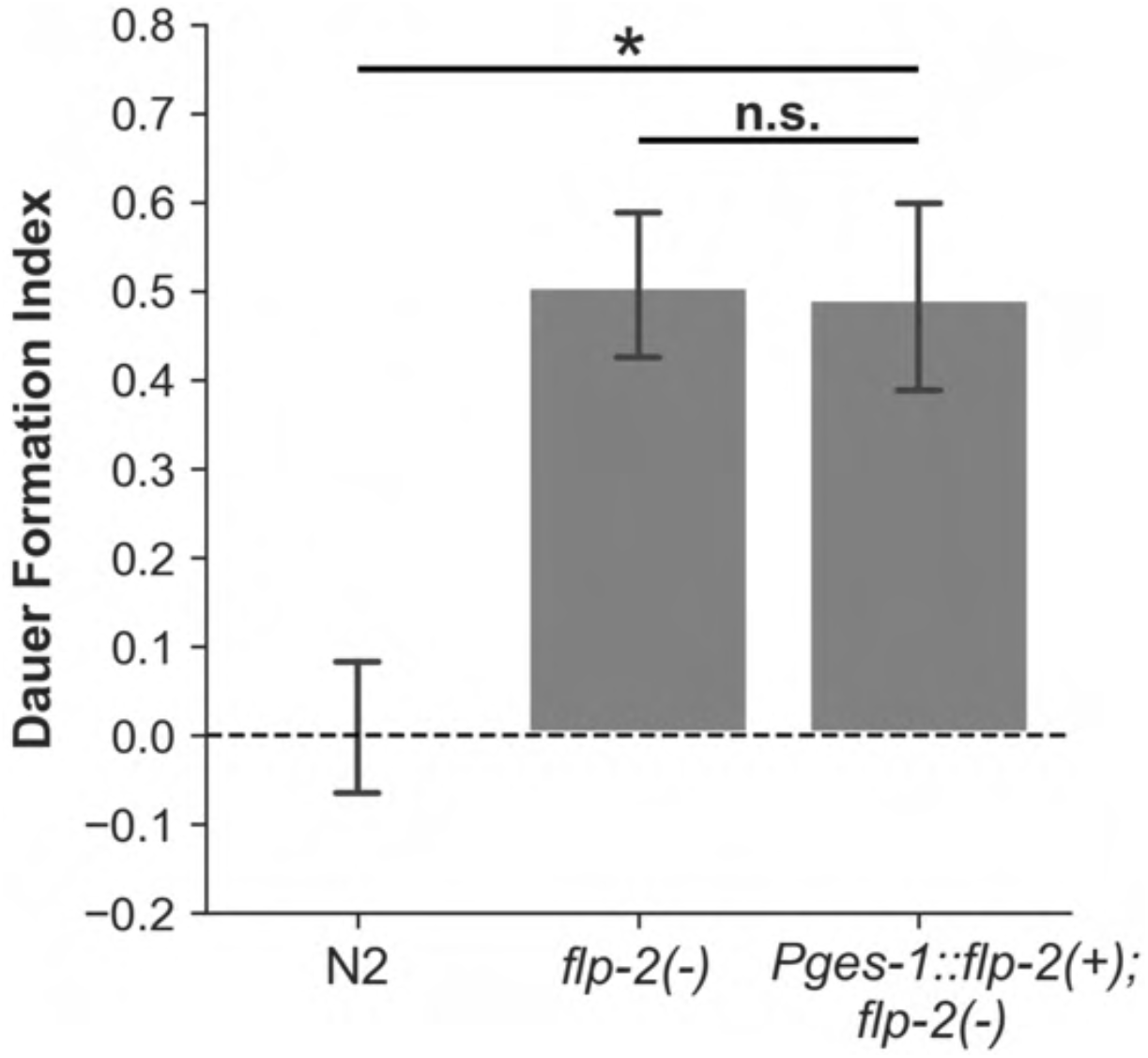
Intestine-specific *flp-2* cDNA expression does not rescue *flp-2(-)* mutant increased dauer formation phenotype. Dauer formation assays for intestine-specific *flp-2* cDNA expression in an *flp-2(-)* mutant background. N=3-4 population assays. Data represented as mean ± SEM, ANOVA followed by Tukey HSD posthoc, *p<0.05, n.s.=no significance.

**Supplementary Figure 2.**
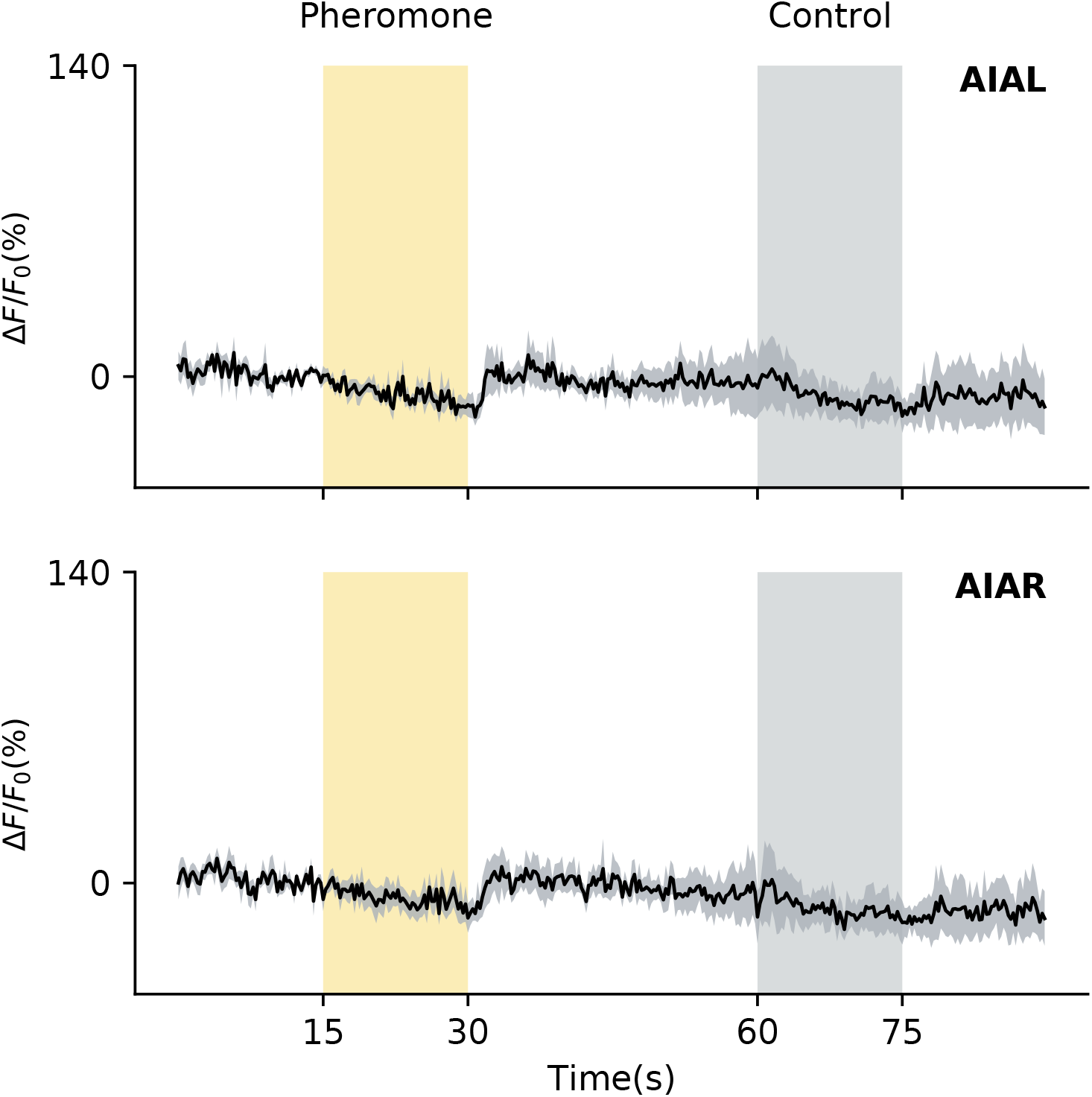
AIA soma GCaMP response to pheromone delivery. *AIA::GCaMP6s* somatic average traces in response to pheromone delivery in microfluidics device. L4 stage animals. Top panel – AIA (Left paired partner). Bottom panel – AIA (Right paired partner). N=5 animals. Data represented as mean (thick black line) ± SEM (gray shading).

**Supplementary Figure 3.**
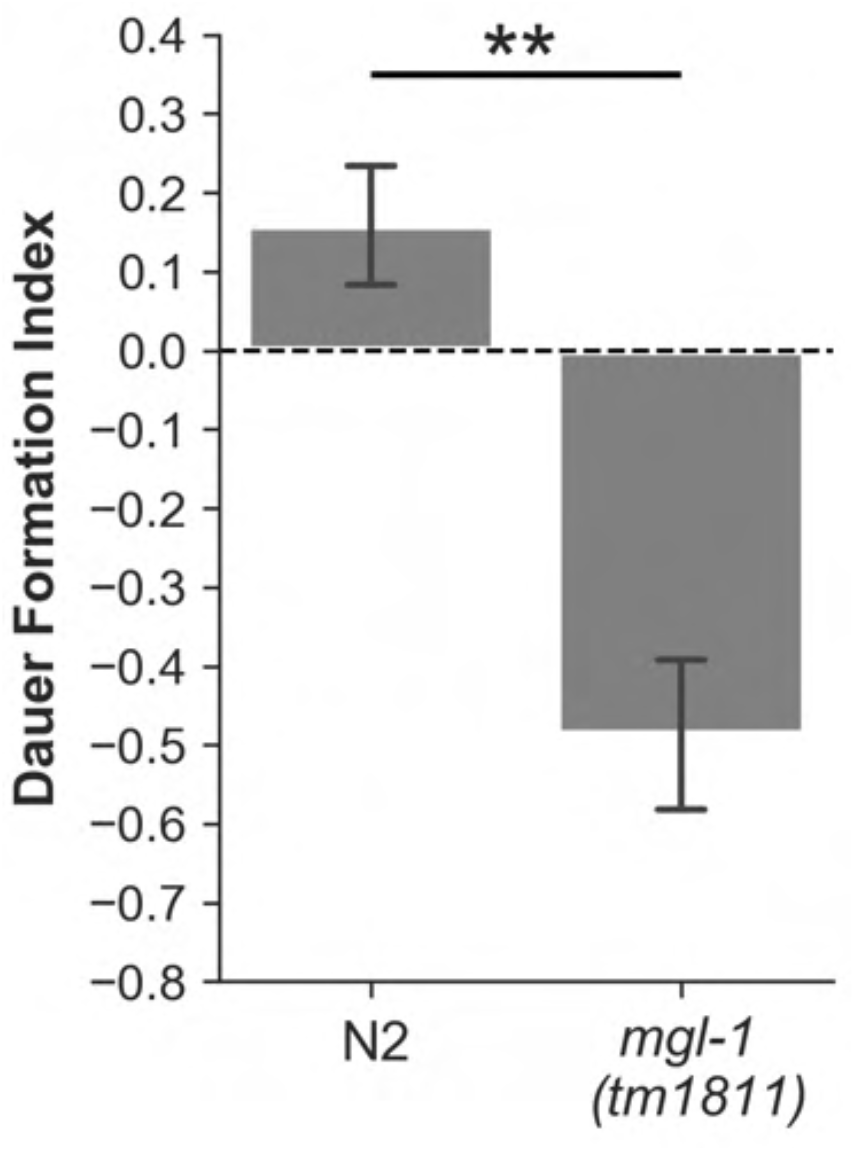
*mgl-1(tm1811)* deletion allele has a decreased dauer formation phenotype. Dauer formation assays for *mgl-1 (tm1811)* mutants. N=4 population assays. Data represented as mean ± SEM, Welch’s T-test, **p<0.01.

**Supplementary Figure 4.**
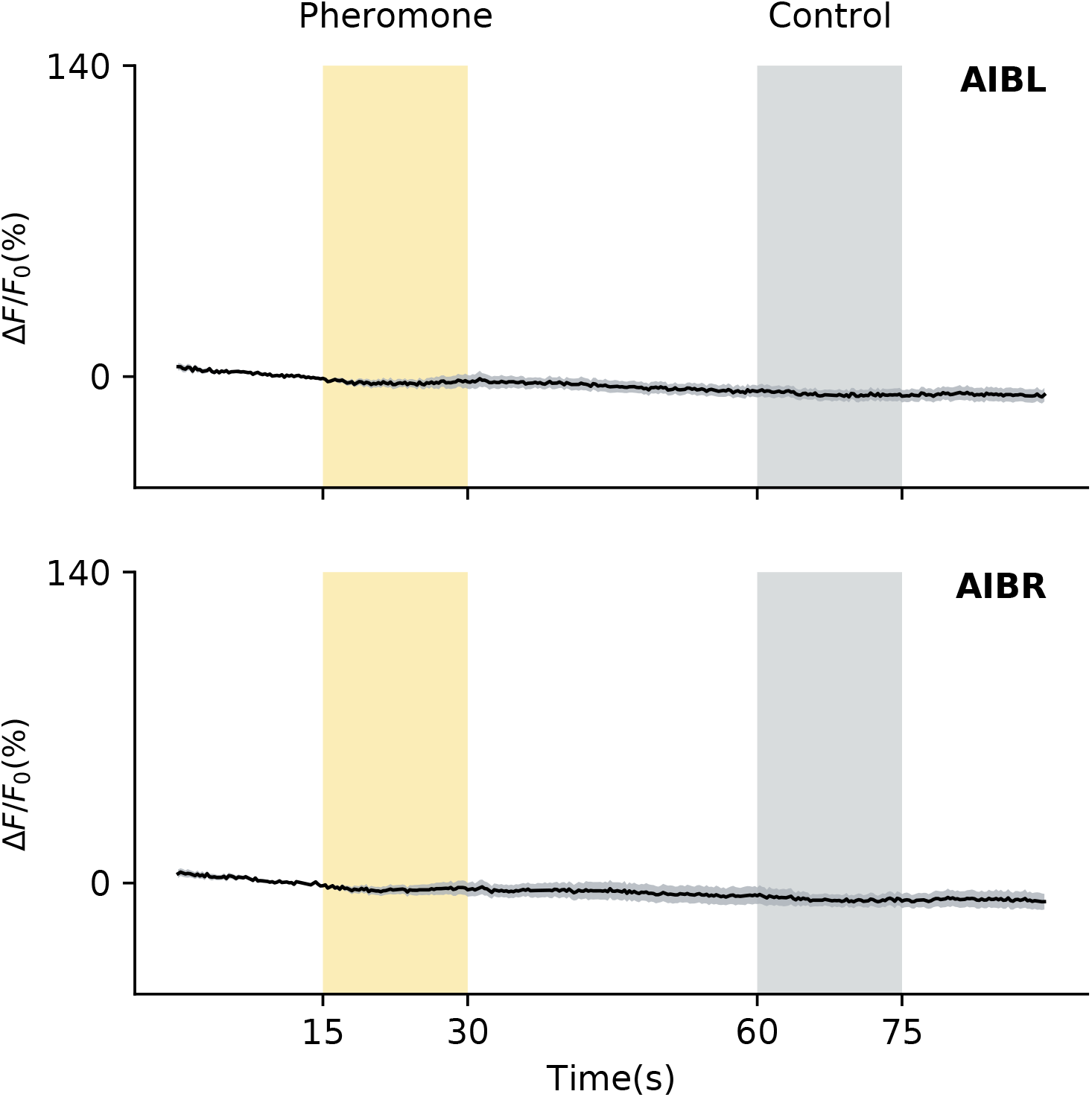
AIB soma GCaMP response to pheromone delivery. AIB::*GCaMP6s* somatic average traces in response to pheromone delivery in microfluidics device. L4 stage animals. Top panel – AIB (Left paired partner). Bottom panel – AIB (Right paired partner). N=13 animals. Data represented as mean (thick black line) ± SEM (gray shading).

## MATERIALS AND METHODS

### Animal maintenance and strains

Animals were cultivated at 21°C on standard nematode growth media (NGM) plates seeded with *Escherichia coli* OP50 cultured in Luria-Bertani (LB) broth. The following strains were used in this study:

**N2** (Bristol)

**PS8919**: *syEx1777[Pflp-2::GFP, Pgcy-28.d::mCherry-H2B, Pofm-1::RFP]*

**PS9101**: *syIs759[Pgcy-28.d::HisCl1::SL2::GFP, Pflp-2::flp-2 cDNA::mCherry, Pofm-1::GFP]*

**PS7370**: *flp-2(ok3351)*

**PS8968**: *flp-2(ok3351); syEx1755[Pflp-2::flp-2* cDNA::SL2::*GFP, Pofm-1::RFP]*

**PS9150**: *flp-2(ok3351); syEx1866[Pges-1::flp-2* cDNA::SL2::*GFP, Pofm-1::RFP]*

**PS8735**: *syEx1755[Pflp-2::flp-2* cDNA::SL2::*GFP, Pofm-1::RFP]*

**PS7866**: *syIs493[Psra-9::cGAL, Pofm-1::RFP]; syIs371[UAS::HisCl1::SL2::GFP, Pofm-1::GFP]*

**PS8159**: *syIs540[Pnpr-9::cGAL, Pofm-1::RFP]; syIs371[UAS::HisCl1::SL2::GFP, Pofm-1::GFP]*

**PS9006**: *syEx1811[Psre-1::HisCl1::SL2::GFP, Pofm-1::RFP]*

**PS8955**: *syEx1793[Psra-9::GCaMP6s, Psre-1::GCaMP6s, Pgcy-28.d::GCaMP6s]*

**PS9111**: *syIs761[Pgcy-28.d::GCaMP6s, Pmyo-2:: mCherry]*

**PS8162**: *syIs543[Pnpr-9::GCaMP6s, Pofm-1::RFP]*

**RB594**: *glc-3(ok321)*

**PS8586**: *npr-9*(*sy1417*)

**PS8628:** *flp-2(ok3351); npr-9(sy1429)*

**PS9192:** *mgl-1(sy1623)*

**PS9193**: *flp-2(ok3351); mgl-1(sy1624)*

**FX01811**: *mgl-1(tm1811)*

**PS7624:** *frpr-18(ok2698)*

**PS7831**: *flp-2(ok3351); frpr-18(ok2698)*

**FX06617**: *npr-30(tm6617)*

**PS8931**: *npr-30(tm6617); syEx1755[Pflp-2::flp-2* cDNA::SL2::*GFP, Pofm-1::RFP]*

**FX01888:** *ins-1(tm1888)*

**PS9137:** *flp-2(ok3351); ins-1(tm1888)*

**PS9147**: *ins-1*(*tm1888*); *syEx1863[Pgcy-28.d::ins-1* cDNA::SL2::*mCherry-H2B*, *Pofm-1::RFP]*

**PS9109**: *npr-30(tm6617); syEx1844[Pnpr-30::npr-30* genomic DNA::SL2::*GFP, Ptax-4::mCherry-H2B, Pofm-1::RFP]*

**PS9152**: *npr-30(tm6617); syEx1868[Pnpr-30::npr-30* genomic DNA::SL2::*GFP, Pgcy-28.d::mCherry-H2B, Pofm-1::RFP]*

### Molecular biology and transgenesis

*flp-2 and ins-1* cDNA were PCR amplified from N2 cDNA library. Promoters and *npr-30* genomic DNA were PCR amplified from N2 genomic DNA library. *Drosophila HisCl1* was PCR amplified from pJL046 (Wang et al., 2017). Constructs were inserted into pSM *GFP*, pSM *mCherry-H2B*, or pSM *GCaMP6s* vector backbone using HiFi Assembly or restriction cloning. *ASK>HisCl1* and *AIB>HisCl1* neuronal silencing strains were generated by placing a *15xUAS::HisCl1::SL2::GFP* effector under the control of ASK and AIB-specific drivers respectively (Wang et al., 2017).

Transgenic strains were generated by microinjection of plasmids with co-injection markers into adult worms. Plasmids were injected at 10-50 ng/μL. Co-injection markers were injected at: *Pmyo-2::mCherry* (0.5 ng/μL), *Pofm-1::RFP* (40 ng/μL), *Pofm-1::GFP* (30-40 ng/μL). 1 kb DNA ladder was used as a filler to bring final DNA concentration to 200 ng/μL.

### CRISPR mutagenesis

CRISPR mutagenesis was performed as described in previously described(Wang et al., 2018). Briefly, a 43-base-pair universal STOP-IN cassette was inserted near the 5’ end of the target gene to disrupt translation. Independent F2 homozygous animals were isolated using PCR detection and confirmed with Sanger sequencing.

### Pheromone-induced dauer formation assays

Crude dauer pheromone extract was prepared as previously described (Schroeder and Flatt, 2014). To control for differences in pheromone potency between batches, a pheromone dose response curve was generated for each batch and the pheromone volume that yielded 50% dauer formation under the experimental conditions described below was selected for dauer formation assays. 8% (w/v) heat-killed OP50 was made by washing bacteria from an overnight culture with virgin S-basal, resuspending with virgin S-basal, and killing at 95-100°C. The afternoon before experiments, L4 worms were picked from well-fed stock plates onto seeded plates and grown overnight at 21°C. Pheromone of desired volume was placed at the center of 35 mm Petri dishes and 2 mL of NGM agar without peptone was poured into each plate. Plates were dried overnight on benchtop at 21°C. On the day of experiments, pheromone plates were seeded with 2 μL heat-killed OP50 and adults animals were allowed to lay 60-70 eggs per plate. Adults were then picked off and 18 μL of heat-killed OP50 added to the plates. Once the food patch had completely dried, plates were sealed with parafilm and placed in a 25.5°C incubator for 72 hours. The number of dauers and non-dauers per plate were counted and the Dauer Formation Index (DFI) per plate calculated using the following formula:

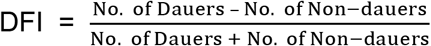

To make histamine plates, histamine dihydrochloride powder was mixed into NGM agar at ~50°C to a final concentration of 10 mM and poured into Petri dishes with pheromone.

For dauer formation assays with diacetyl, assays were set up as described above. Before parafilming, 2 μL of 200 proof ethanol or 2 μL or 1:1000 diacetyl diluted in ethanol was added to the side of the food patch.

For experiments using extrachromosomal array transgenic lines, lines that had 70-80% array transmission rates were selected for population assays. All animals on each plate were counted unless noted otherwise. At least 3 population assays were completed for each control and genotype/treatment (except for N2 controls in Figure 7B), with each assay containing 60-70 animals. Control and genotype/treatment plates were always assayed simultaneously under the same conditions for comparisons.

### Microscopy

Animals were immobilized with 50 mM sodium azide on 2% agarose pads. Slides were imaged under a ZEISS Imager.Z2 microscope attached to a Axiocam 506 mono camera capture source. Images were processed using ZenPro software.

### Calcium imaging

#### Microfluidic device fabrication

We designed a 2-layer microfluidic chip capable of delivering sequences of stimuli with a worm trap suitable for housing worms at L4 larval stage. The chip was designed in AutoCAD software, and sent to CAD/Art Services, Inc to print the photomask. Photolithography in a clean room was performed on a silicon wafer to make the 2-layer mold from the photomask. For the first layer, which included the worm trap, SU-8 2025 was spin coated on the silicon wafer at 4000 rpm to achieve 25 μm thickness. For the second layer, the same photoresist was spin coated at 1250 rpm for a thickness of 70 μm. Polydimethylsiloxane (PDMS) was poured over the mold and cured on a 90°C hotplate to solidify. Each PDMS chip was then punched with a 1 mm biopsy punch and was bonded to a cover slip using a handheld corona treater.

#### Microfluidic experiment design

L4 stage animals were assayed and all experiments were conducted at 20°C. We chose 3 stimuli for the experiment: 10% (v/v) crude pheromone extract diluted in CTX buffer (5 mM KH_2_PO_4_/K_2_HPO_4_ pH 6, 1 mM CaCl_2_ and 1 mM MgSO_4_,50 mM NaCl) as the main stimulus, CTX buffer as the control stimulus, and 0.01 mM Fluorescein to confirm chemical delivery. Each stimulus interval was 15 seconds, sufficient time to capture the rise and fall of calcium transients. Stimulus interval was then followed by a 30-second CTX buffer interval to give neurons enough time to return to baseline and be ready for the next stimulus.

#### Data Analysis

For each animal, fluorescence was captured at 4 volumes/second, 25 z-slices/volume and 1.2 μm/slice and average background intensity was subtracted from each volume. To measure calcium activity of the neuron’s soma, average intensity of pixels within a 3.2 μm x 3.2 μm x 3.6 μm volume around the center of the soma was calculated. To extract calcium activity of neuron processes, a 38 μm x 38 μm x 24 μm rectangular volume surrounding the processes were identified at each timepoint, pixels from the soma were removed from the volume and the average intensity of the remaining pixels were calculated. F_0_ was defined as average intensity during the 5-second window before the delivery of pheromone.

## STATISTICAL ANALYSIS

For dauer formation assays, unpaired Welch’s T-test or ANOVA followed by Tukey HSD posthoc analysis were used to determine statistically significant differences between groups. All statistical analyses were performed in R. All plots were generated in Python.

